# Xenotransplanted human cortical neurons reveal species-specific development and functional integration into mouse visual circuits

**DOI:** 10.1101/626218

**Authors:** Daniele Linaro, Ben Vermaercke, Ryohei Iwata, Arjun Ramaswamy, Brittany A. Davis, Leila Boubakar, Baptiste Libé-Philippot, Angéline Bilheu, Lore De Bruyne, David Gall, Klaus Conzelmann, Vincent Bonin, Pierre Vanderhaeghen

## Abstract

How neural circuits develop in the human brain has remained almost impossible to study at the neuronal level. Here we investigate human cortical neuron development, plasticity and function, using a mouse/human chimera model in which xenotransplanted human cortical pyramidal neurons integrate as single cells into the mouse cortex. Combined neuronal tracing, electrophysiology, and *in vivo* structural and functional imaging revealed that the human neurons develop morphologically and functionally following a prolonged developmental timeline, revealing the cell-intrinsic retention of juvenile properties of cortical neurons as an important mechanism underlying human brain neoteny. Following maturation, human neurons transplanted in the visual cortex display tuned responses to visual stimuli that are similar to those of mouse neurons, indicating capacity for physiological synaptic integration of human neurons in mouse cortical circuits. These findings provide new insights into human neuronal development, and open novel experimental avenues for the study of human neuronal function and diseases.

**Highlights:**

- Coordinated morphological and functional maturation of ESC-derived human cortical neurons transplanted in the mouse cortex.
- Transplanted neurons display prolonged juvenile features indicative of intrinsic species-specific neoteny.
- Transplanted neurons develop elaborate dendritic arbors, stable spine patterns and long-term synaptic plasticity.
- In the visual cortex transplanted neurons display tuned visual responses that resemble those of the host cortical neurons.

## Introduction

The brain and cerebral neocortex in particular increased rapidly in size and complexity during human evolution (Dehay et al., 2015; Geschwind and Rakic, 2013; Gotz and Huttner, 2005; Lui et al., 2011). While there has been major progress in understanding the mechanisms underlying the increase in human cortical size (Borrell and Gotz, 2014; Florio et al., 2015; Hansen et al., 2010; Stahl et al., 2013; Suzuki et al., 2018), this only explains a part of our brain evolution: features that are thought to be unique to human brain development mostly lie in the way neurons assemble to form functional and complex neural circuits (Bae et al., 2015; Defelipe, 2011; Geschwind and Rakic, 2013; Sousa et al., 2017).

How human neurons assemble into mature circuits has important implications for brain evolution and diseases. On the one hand, a remarkable feature of the human cortical circuits, compared with non-human species, is their unusually prolonged development (Petanjek et al., 2011; Rakic et al., 1986). This neoteny, or retention of juvenile traits in a more mature organism, has long been proposed to play a key role in the acquisition of human-specific cognitive features (Gould, 1992), and is thought to be mostly the consequence of a prolonged pattern of maturation of cortical neurons compared with non-human primates (Bufill et al., 2011; Defelipe, 2011; Petanjek et al., 2011). While cortical neurons maturation takes 3-4 weeks in the mouse and 3-4 months in the macaque, neurons of the human cortex mature over years postnatally. Compared with macaque neocortex, human cortical pyramidal neurons display prolonged periods of dendritic outgrowth and branching, as well as dendritic spine and synapse formation (over months to several years), and synaptic pruning (up to a decade) (Huttenlocher, 1979; Huttenlocher et al., 1982a, b; Mrzljak et al., 1990; Petanjek et al., 2011), while comparison with chimpanzee also indicates human neoteny though less pronounced (Bianchi et al., 2013; Liu et al., 2012). Beyond evolution, human neuron development has important implications to understand brain diseases (Liu et al., 2016). Indeed, alterations of neuronal maturation have been strongly associated with cognitive dysfunction, in particular intellectual disability (ID) and autism spectrum disorders (ASD), (Hutsler and Zhang, 2010; Pfeiffer et al., 2010; Takashima et al., 1981; Willsey et al., 2013), although the underlying pathological mechanisms remain mostly unexplored in human neurons.

Despite their importance to understand human brain evolution and diseases, our basic knowledge about human neuron and synapse development has remained fragmentary. This is mostly because of the difficulty of studying live developing human neurons, especially in the context of a brain circuit. Pluripotent stem cell (PSC)-based models offer new opportunities to study human neural development (Astick and Vanderhaeghen, 2018; Di Lullo and Kriegstein, 2017; Suzuki and Vanderhaeghen, 2015; Zeltner and Studer, 2015). However they have remained difficult to use to study cortical neuron development *in vitro*, given the technical challenges of keeping them in vitro for months-long periods (Di Lullo and Kriegstein, 2017; van den Ameele et al., 2014). Another approach being developed is xenotransplantation of PSC-derived human neurons into the mouse brain, which in principle can provide a mean to study long-term human neuronal maturation under more physiologically realistic conditions. Human PSC-derived pyramidal cortical neurons transplanted in the mouse brain develop morphologically and synaptically within the host mouse brain (Espuny-camacho et al., 2013), and interestingly, transplanted human neurons develop more slowly than similarly transplanted mouse neurons (Gaspard et al., 2008). However, it has remained unclear whether xenotransplanted neurons, although competent to make synapses and receive inputs (Espuny-camacho et al., 2013; Espuny-Camacho et al., 2018; Mansour et al., 2018; Qi et al., 2017; Real et al., 2018; Tornero et al., 2017), can actually integrate physiologically to participate to the function of cortical circuits. Notably in this context, to date xenotransplantation studies focused on cell-compact transplants that comprise a large number of cells lumped together with limited access to the host tissue (Espuny-camacho et al., 2013; Espuny-Camacho et al., 2018; Mansour et al., 2018; Real et al., 2018; Tornero et al., 2017). While these transplants are viable and can host vascular and glial cells, they form a local environment that differs dramatically from the host cortical tissue, and may not provide all necessary cues for proper circuit integration. A recent study using in vivo imaging showed indeed limited synaptic maturation and integration of human neurons in mouse cortex (Real et al., 2018).

Therefore, whether xenotransplanted human neurons can successfully mature, integrate and gain information processing function is not known. Likewise, it remains unknown whether the prolonged neuronal development observed for transplanted human cortical neurons actually reflects these partially isolated graft conditions, or is the genuine result of an intrinsic developmental programme physiologically relevant to human brain neoteny.

Here we developed a novel experimental model to address these questions and explore human cortical neuronal development *in vivo*, using xenotransplanted human cortical pyramidal neurons that integrate as single cells into the neonatal mouse cortex. We show that these human cortical neurons develop following a coordinated roadmap that recapitulates the key milestones of cortical neuronal development in the human brain, indicating that human cortical neuronal neoteny has a strong intrinsic component. Moreover following maturation, the neurons become highly connected with the host mouse cortical neurons, and display responses to sensory stimuli that resemble those of host neurons, thus providing novel experimental tools to study human neuronal function *in vivo*.

## Results

### Intraventricular human cortical xenotransplantation leads to robust integration in the mouse cortex

A major limitation of current xenotransplantation approaches to study cortical development is that transplanted cells tend to stick to each other and do not integrate well into the host brain tissue. To circumvent this technical hurdle, we developed a transplantation procedure that increases the rate of integration of human cortical pyramidal neurons into the neonatal cortex. Specifically we took advantage of the observation that EGTA, added together with the cells during intraventricular injection, can promote integration and migration of mouse and human neurons into the mouse host cortex (Nagashima et al., 2014) (Figure 1A, S1A). In these conditions, transplanted human ESC-derived cortical pyramidal neurons, examined 2-8 weeks post-transplantation, were found within the cortical gray matter, from deep (5/6) to superficial (2/3) cortical layers, displaying a radial orientation (Figure 1B, S1B) and expressing markers (Figure 1C-D, S1C-D) typical of cortical pyramidal neurons (Greig et al., 2013). Transsynaptic rabies experiments were performed to probe the presynaptic partners of human neurons, by transplanting human neurons transduced with TVA-mCherry fusion and rabies glycoprotein, followed by GFP-expressing rabies virus intracortical injection. This revealed that at 4 months post-transplantation (MPT) human neurons receive abundant connections from mouse cortical neurons (Figure 1E).

**Figure 1.**
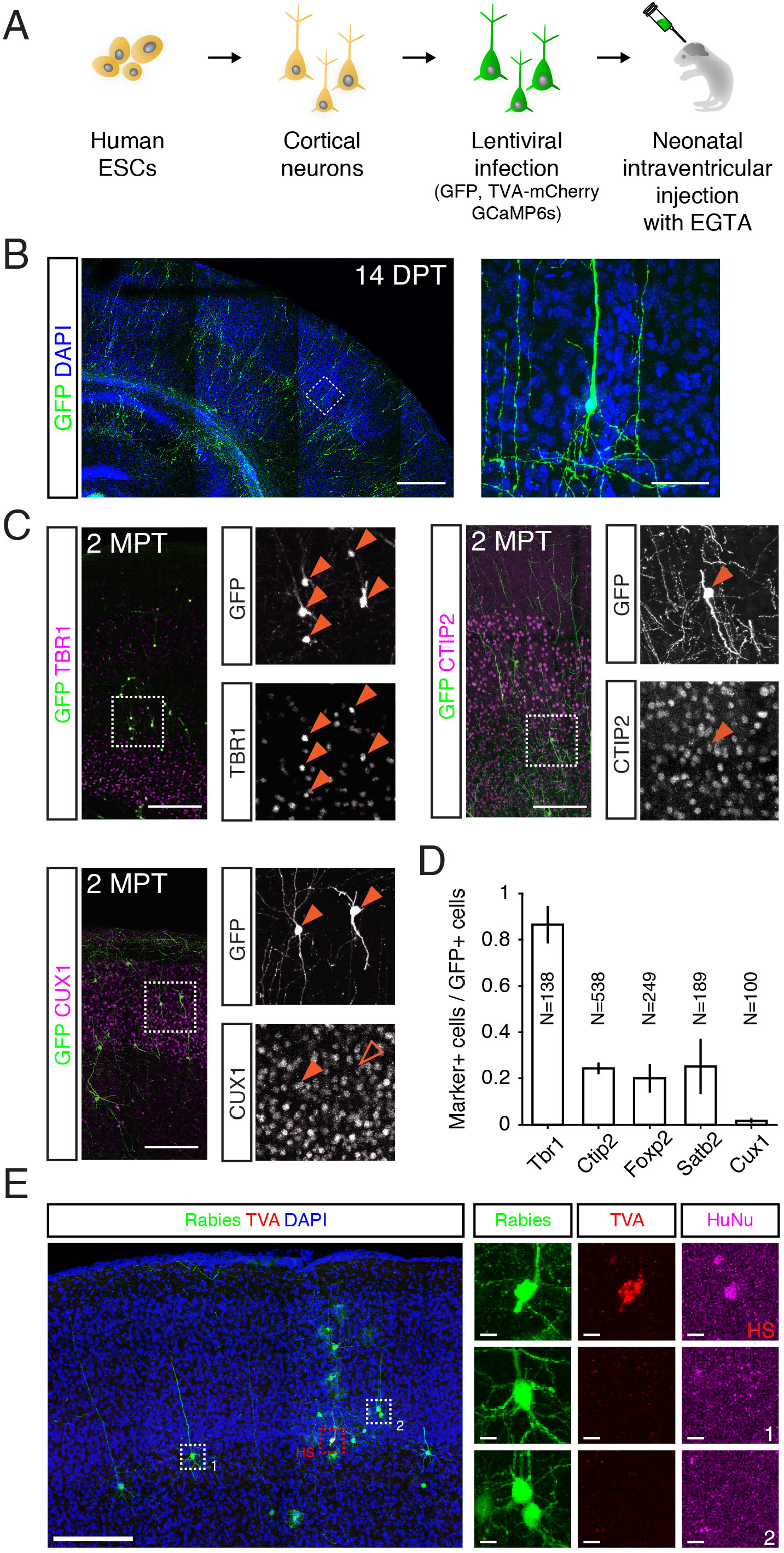
Transplanted hESC-derived cortical neurons integrate into the mouse cortex and establish synaptic connections with the host tissue. (A) Scheme representing differentiation and transplantation protocol. (B) Left: representative low-magnification image of hESC-derived cortical neurons transplanted at birth into the mouse cortex, 14 days post-transplantation. Right: image area highlighted by the dashed box in the left image. Notice the radial orientation and the apical processes of the GFP+ human cells. (C) Examples of isolated human cells positive for deep- and upper-layer cortical markers. (D) Proportion of human cells positive for several cortical markers. *N* refers to the total number of counted GFP+ human cells. (E) Left: representative low-magnification image of starter transplanted hESC-derived cortical neurons and presynaptic mouse cells. Right: high-magnification images of the areas highlighted with dashed boxes on the left. HS indicates human starter cell (positive for TVA-RFP and human nucleus (hunu) markers, and infected with GFP-expressing rabies virus). The other rows show two examples of presynaptic mouse pyramidal cells, infected with rabies virus (GFP+, HuNu-, TVA-FRP-). Scale bars: (B) left, 500 *µ*m, right, 50 *µ*m. (C) 200 *µ*m. (E) Low magnification image, 200 *µ*m, high magnification zooms, 10 *µ*m.

### Coordinated functional and morphological neotenic development of human cortical neurons

To characterize the maturation of xenotransplanted human neurons in the mouse cortex, we first performed ex vivo whole-cell patch-clamp recordings and measured the progression of their action potential shape, firing patterns, intrinsic and extrinsic properties from 1 to 11 MPT (Figure 2, Figure S2, Table S1). Transplanted human neurons showed progressively maturing firing patterns, from low amplitude action potentials and rapidly adapting responses at 1 MPT to sustained firing and mature-looking spikes at 10 MPT (Figure 2A), that are virtually indistinguishable from those recorded in adult rodents (Jiang et al., 2015) and humans (Beaulieu-Laroche et al., 2018; Testa-Silva et al., 2014). These changes in firing patterns were accompanied by changes in intrinsic properties, including progressive membrane potential hyperpolarization (Figure 2B), a decrease in input resistance of approximately one order of magnitude (Figure 2C, Figure S2C), a substantial increase in maximum sodium currents (Figure 2D) and a shortening of AP half-width with an associated increase in spike amplitude (Figure S2A-B). These changes were associated with a concomitant decrease of spontaneous firing (Figure 2F,G), in line with the membrane potential hyperpolarization observed over the same developmental period, and an increased capacity to fire at high rates (Figure 2E), indicative of progressive maturation and increase in density of voltage-dependent ion channels present in the cell membrane. Importantly, recordings of spontaneous excitatory post-synaptic currents showed a progressive increase in the rate of incoming spontaneous synaptic events over the course of several months (Figure 2H), indicating synaptic integration within the host cortical tissue.

**Figure 2.**
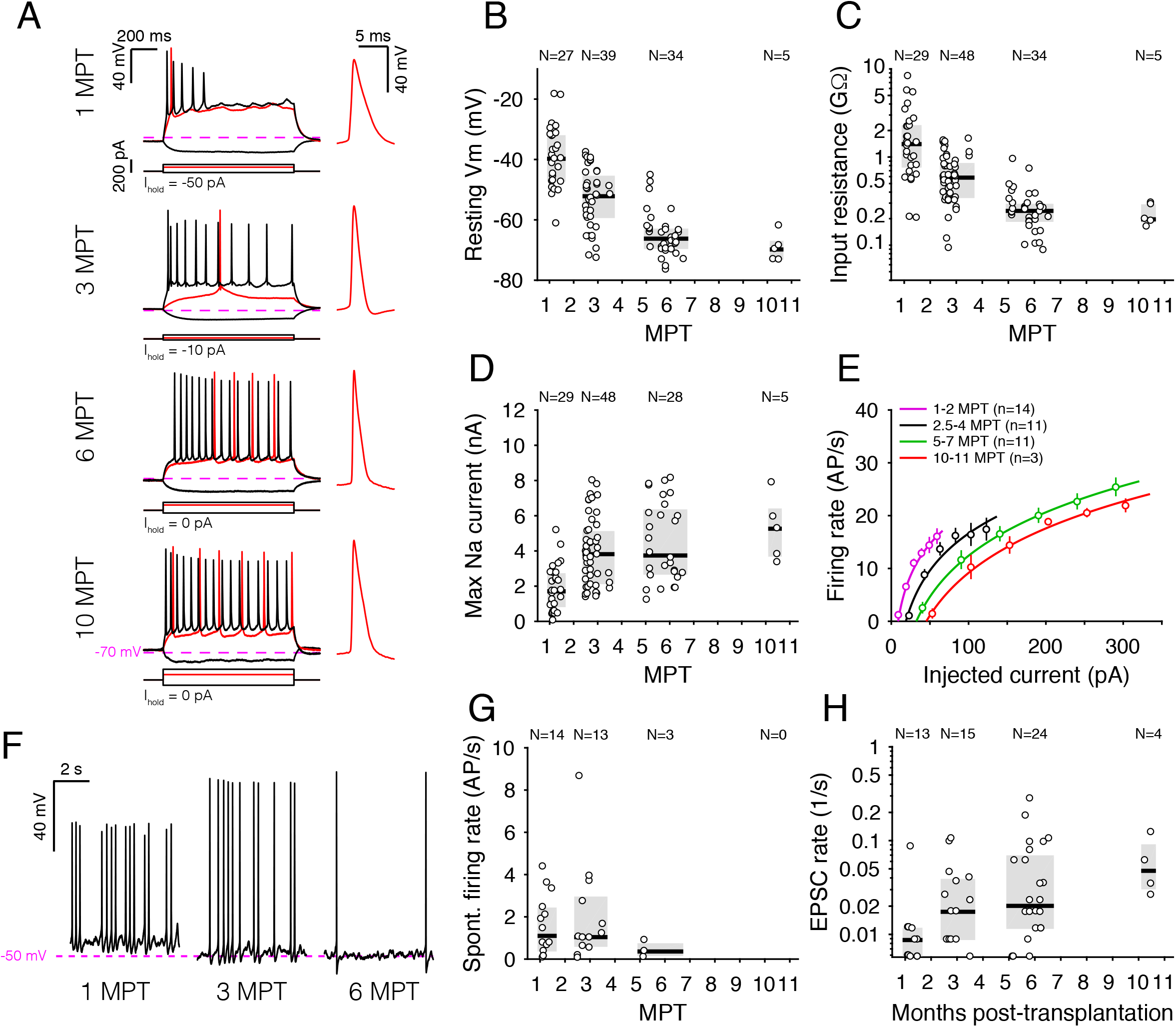
Electrophysiological properties of hESC-derived cortical neurons over the course of several months. (A) Membrane potential dynamics in response to the injection of hyperpolarizing and depolarizing steps of current at various stages of development. Inset: first AP emitted upon injection of current around the rheobase amplitude. Dashed magenta line indicates the value of −70 mV, at which cells were kept by the injection of the offset current I_hold_ indicated in each panel. (B-D) Population data of several electrophysiological quantities over the course of 11 MPT. Notice how some properties (resting Vm and maximum sodium current) appear to be still evolving at 11 MPT, while others (input resistance and AP half-width) seem to have reached a plateau level. In each panel, N indicates the number of cells for which a given feature was recorded. (E) Static input-output characterization (f-I curves) of cells grouped based on their age. Markers and error bars indicate mean ± SEM. Continuous lines are fits to the data with a power-law function of the form *f*(*I*) = *aI*^*b*^ + *c*. (F) Representative examples of spontaneously active cells (I_hold_ = 0 pA). (G) Population summary of spontaneous firing rates as a function of age: no cells displayed spontaneous firing rate later than 6 MPT. (H) Rate of spontaneous excitatory post-synaptic currents as a function of age. Pooled data in B, C, D, G and H are represented as median and interquartile range.

Collectively these data indicate that transplanted human neurons show progressive maturation in intrinsic and extrinsic properties at least until 11 MPT, at which time-points most of the values measured are comparable to those of human or mouse neurons at adult stages (Beaulieu-Laroche et al., 2018; Testa-Silva et al., 2014). This is in stark contrast with the developmental pace of mouse neurons transplanted in the mouse cortex, which takes place in only a few weeks (Falkner et al., 2016; Gaspard et al., 2008; Michelsen et al., 2015). Notably, some of the cortical neurons analyzed were generated using another protocol of in vitro corticogenesis (Shi et al., 2012), and their functional features displayed highly similar timing of maturation (Figure S2D-E). This further suggests that this is a robust and general feature of human cortical neurons in this transplantation paradigm, which occurs independently from the conditions of in vitro differentiation.

A fundamental feature of cellular development is that it follows a highly coordinated pattern, both morphologically and functionally. To explore this important aspect, a subset of the recorded neurons (N=28) were filled with biocytin and their morphologies digitally reconstructed and analyzed. We observed over time (1-7MPT) pronounced morphological changes both in terms of dendritic length and complexity (Figure 3A-C, Figure S3A-C) as well as dendritic spine density and morphology (Figure 3D-F). Morphological and functional parameters correlated strongly with each other, including for instance resting membrane potential with dendritic length and spine density (Figure 3G,H).

**Figure 3.**
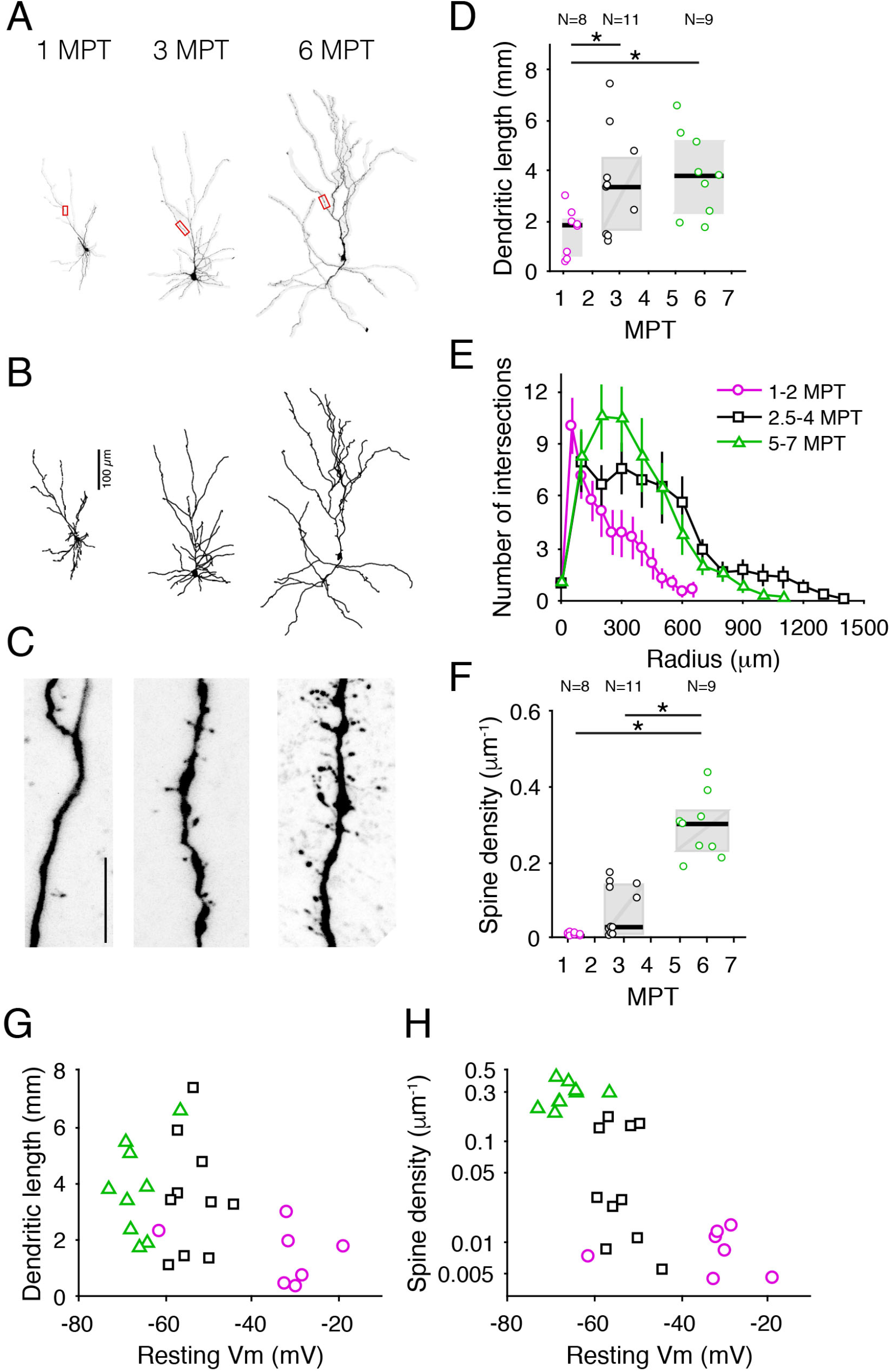
Morphological properties of hESC-derived cortical neurons over the course of several months. (A) Confocal images of representative neurons at three successive developmental time points. (B) Digital reconstructions of the cells shown in A. (C) High magnification images of the dendritic branches highlighted in red in A: notice the appearance of spines at around 3 MPT and the significant increase in spine density at around 5 MPT. Scale bar, 10 *µ*m. (D) Dendritic length as a function of age for 28 digitally reconstructed cells. (E) Sholl analysis for the reconstructed cells, segregated in 3 groups according to their age. Each symbol represents a cell, magenta circles are cells aged 1-2 MPT, black squares are cells aged 2.5-4 MPT and green triangles are cells aged 5-7 MPT. Markers and error bars indicate mean ± SEM. (F) Spine density as a function of age. Notice the marked separation between cells before 4 MPT and after 5 MPT. (G-H) Dendritic length and spine density as a function of resting Vm. Markers and colors as in E. In D and F, pooled data are represented as median and interquartile range (*p < 0.05, one-way ANOVA).

Taken together, these electrophysiological and morphological data indicate that xenotransplanted human cortical neurons follow a coordinated program of functional and morphological maturation following a time-line that is strikingly similar to the one observed in the human cortex in vivo, despite the fact that all the analyzed neurons developed as isolated cells within the much faster-developing mouse cortical tissue. These data therefore strongly suggest that this developmental programme – in particular the neotenic-like prolonged time-line - is largely intrinsic to the human neurons.

### Transplanted human neurons display rich dendritic spine structural dynamics in adult mouse cortex

As transplanted human neurons develop according to a prolonged timeline, they retain juvenile properties in an otherwise mature mouse brain. Another salient feature of juvenile neurons is their structural dynamics, which precedes the stable establishment of synaptic connectivity (Grutzendler et al., 2002; Zuo et al., 2005). A previous study showed very low spine survival rates of human neurons in the mouse cortex with little signs of stabilization (Real et al., 2018). To assess dynamic aspects of synaptic maturation and cellular integration of transplanted developing neurons, we performed chronic longitudinal cellular imaging of dendritic spines in the mouse cortex (Figure 4). Xenotransplanted animals (n=5) were imaged using a two-photon microscope through a chronically-implanted glass window positioned over occipital cortex (Figure 4A).

**Figure 4.**
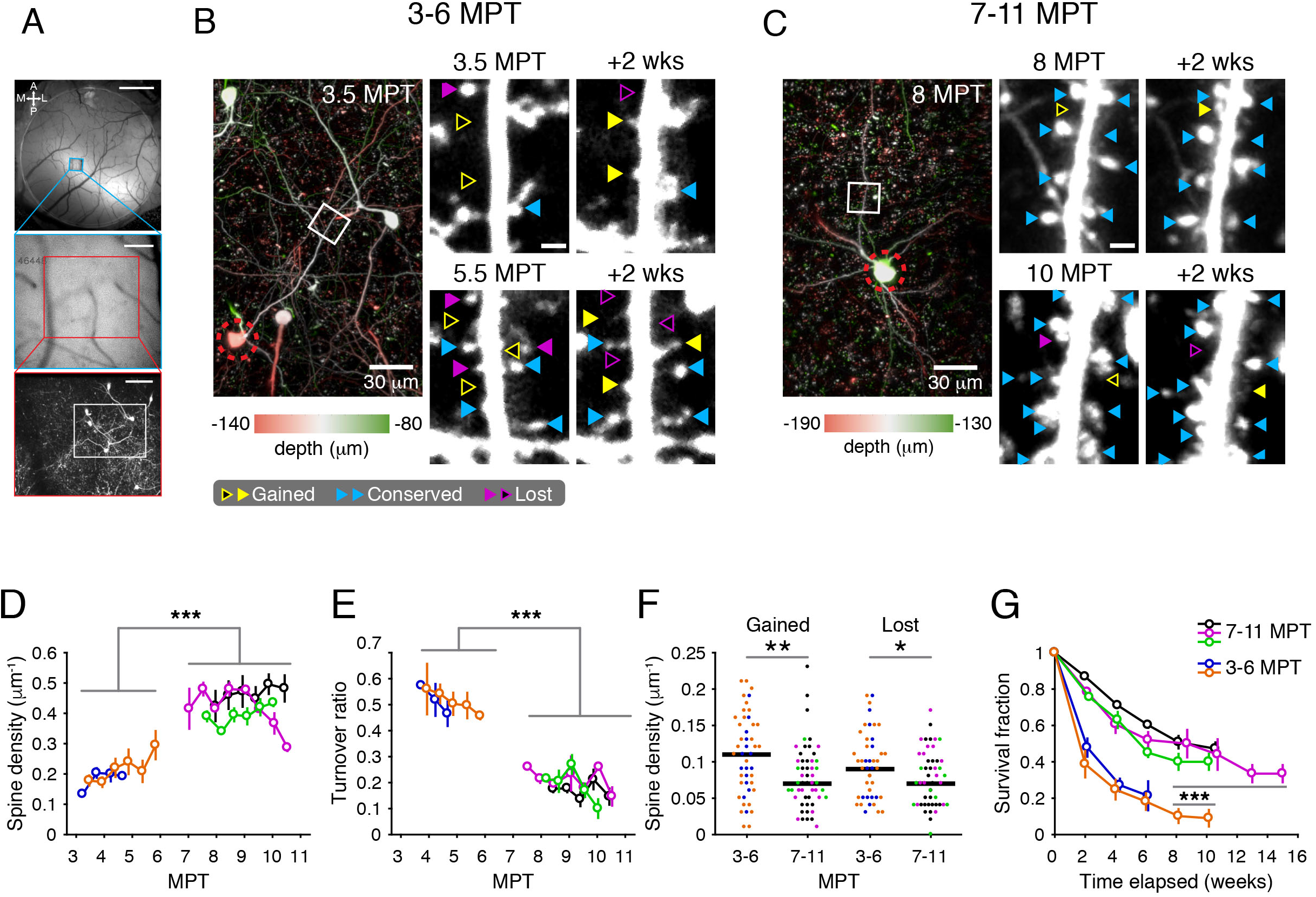
In vivo imaging reveals structural dynamics of dendritic spines of transplanted human neurons. A) Outline of strategy used to relocate cells over time. *Top*: Low magnification wide-field image showing the cranial window at the start of the experiment. The blood vessels serve as landmarks which allow navigation to a specific field-of-view. *Middle*: Higher magnification of region marked with a blue box in *top*. Wide-field image taken with a 25x water immersion objective. At this resolution, we can more precisely match to a location imaged in a previous session. *Bottom*: High magnification 2-photon image of the region marked in red in *middle*. Maximum intensity projection of a 200 *µ*m stack centered around the hESC somata shown in B) - note a 90 deg. clockwise rotation is applied panel B). Scale bars represent top: 1 mm, middle: 200 *µ*m, bottom: 100 *µ*m. (B) Example data showing hESC neurons at 3-6 MPT. *Left*: the soma (red dashed circle) and the branch segment (white box) shown in detail on the right. Color code reflects depth ranging from green (towards pial surface) over white (matches depth of marked branch) to red (towards white matter). *Right*: (rows) same branch segment at 3.5 MPT and 5.5 MPT and their 2 week sequent timepoint (columns). Spines annotations are marked with arrowheads using following color conventions: yellow denotes gained spines, purple indicates lost spines and blue marks conserved spines (open arrowheads indicate locations where spines were/will be present in the adjacent timepoint). (C) Example data showing a hESC neuron at 7-10 MPT. Same conventions as in B). Note the relative increase of blue arrowheads and concurrent decrease of yellow/purple arrowheads. (D) Summary of spine density results. Spine density increases between cells aging 3-6 MPT (n=2 animals) and 7-10 MPT (n=3 animals). Colors indicate animal identity and are consistent over panels D-G. (E) Summary of spine turnover results. Spine turnover decreases between cells aging 3-6 MPT (n=2 animals) and 7-10 MPT (n=3 animals). (F) Comparison of spine gain and loss. Both density of gained and lost spines decreases between cells aging 3-6 MPT (n=2 animals) and 7-10 MPT (n=3 animals).(G) Summary of survival fraction results. Survival fraction increases between cells aging 3-6 MPT (n=2 animals) and 7-10 MPT (n=3 animals).Markers and error bars indicate mean ± SEM. (*p<0.05, **p<0.01, ***p<0.001, ranksum test).

The GFP labeled cells were located using a combination of widefield imaging and 2-photon microscopy (Holtmaat et al., 2006; Trachtenberg et al., 2002; Zuo et al., 2005), and the same cells and dendrites were imaged at 2-week intervals for periods up to 12 weeks, focusing on groups of mice that were 3-6 or 7-11 MPT when we started the timelapse imaging (Figure 4B-C).

These experiments revealed that xenotransplanted human cortical neurons display rich spine dynamics comparable to those observed in developing mouse cortical neurons at juvenile stages, whether native or following transplantation (Cruz-Martin et al., 2010; Falkner et al., 2016). Consistent with cortical slice data (Figure 3F), xenotransplanted human cortical neurons showed increase in spine density for up to 10-12 months post transplantation (Figure 4D; Supplementary Table S1), reaching values that are in line with those reported for human neurons at 1-2 years of age (Jacobs et al., 1997; Jacobs et al., 2001; Petanjek et al., 2011) and in the adult mouse visual cortex (Holtmaat et al. 2005).

Remarkably, the spine turnover ratio, a dynamic measure of cortical dynamics and plasticity (Holtmaat et al., 2005), was still quite high at 3-6 MPT, reflecting juvenile properties of the neurons, but then dropped at 7-10 MPT, indicating that the neurons had reached a more stable pattern of spine dynamics (Figure 4E) although still more unstable than the one found in adult mouse cortex (Grutzendler et al., 2002; Zuo et al., 2005). Indeed this change of spine turnover ratio was in part followed by a pronounced reduction in number of gained and lost spines (Figure 4F; Supplementary Table). By following spines across multiple consecutive imaging sessions, we also observed a flattening of the spine survival function (Figure 4G) reflecting an increase in spine survival rates from ∼2 weeks at 3-6 MPT to ∼8 weeks at 7-11 MPT (>50% of spines surviving; Supplementary Table).

Taken together, these data support the notion that in terms of spine development and dynamics, transplanted human neurons follow a prolonged timeline like native human neurons (Petanjek et al., 2011), that is largely intrinsic to these neurons, as they still display juvenile-like dendritic spine dynamic patterns at 3-6MPT, i.e. in a mature mouse brain environment in which the cortical neurons display mostly stable spine dynamics (Falkner et al., 2016; Holtmaat et al., 2005; Tjia et al., 2017; Zuo et al., 2005). Moreover and importantly, the stabilization of the dendritic spines at later stages support the view that the neurons eventually integrate synaptically with the host brain despite their prolonged timing of maturation.

### Transplanted human neurons display long-term synaptic potentiation

The observation of structural dynamics, together with an increase in the rate of incoming spontaneous synaptic events over the course of several months (Figure 2H), raised the possibility that transplanted human neurons might display synaptic plasticity. To assess this, we applied a long-term potentiation (LTP) protocol (Nevian and Sakmann, 2006) to xenotransplanted neurons in ex vivo slice preparations (Figure 5A). We found that human cells aged 3 to 6 MPT were capable of displaying potentiation of their response, measured as the amplitude of the excitatory post-synaptic potential (EPSP), lasting for over 30 minutes after the application of the pairing stimulus (Figure 5B). At the population level, we found that about two thirds of the tested human neurons displayed LTP (N=7/10, Figure 5C), whereas the remaining third (N=3/10) remained stable throughout the course of the recording (Fig. 5D). We conducted analogous experiments in L5 pyramidal cells in the primary somatosensory cortex of 3 months-old NOD/SCID mice and found that LTP can be reliably induced also in these cells, albeit intriguingly the magnitude of the response increase is smaller in murine cells (Fig. 5E). Taken together, these results indicate that xenotransplanted human cortical neurons present robust functional synaptic plasticity comparable to those observed in the host cortical pyramidal neurons.

**Figure 5.**
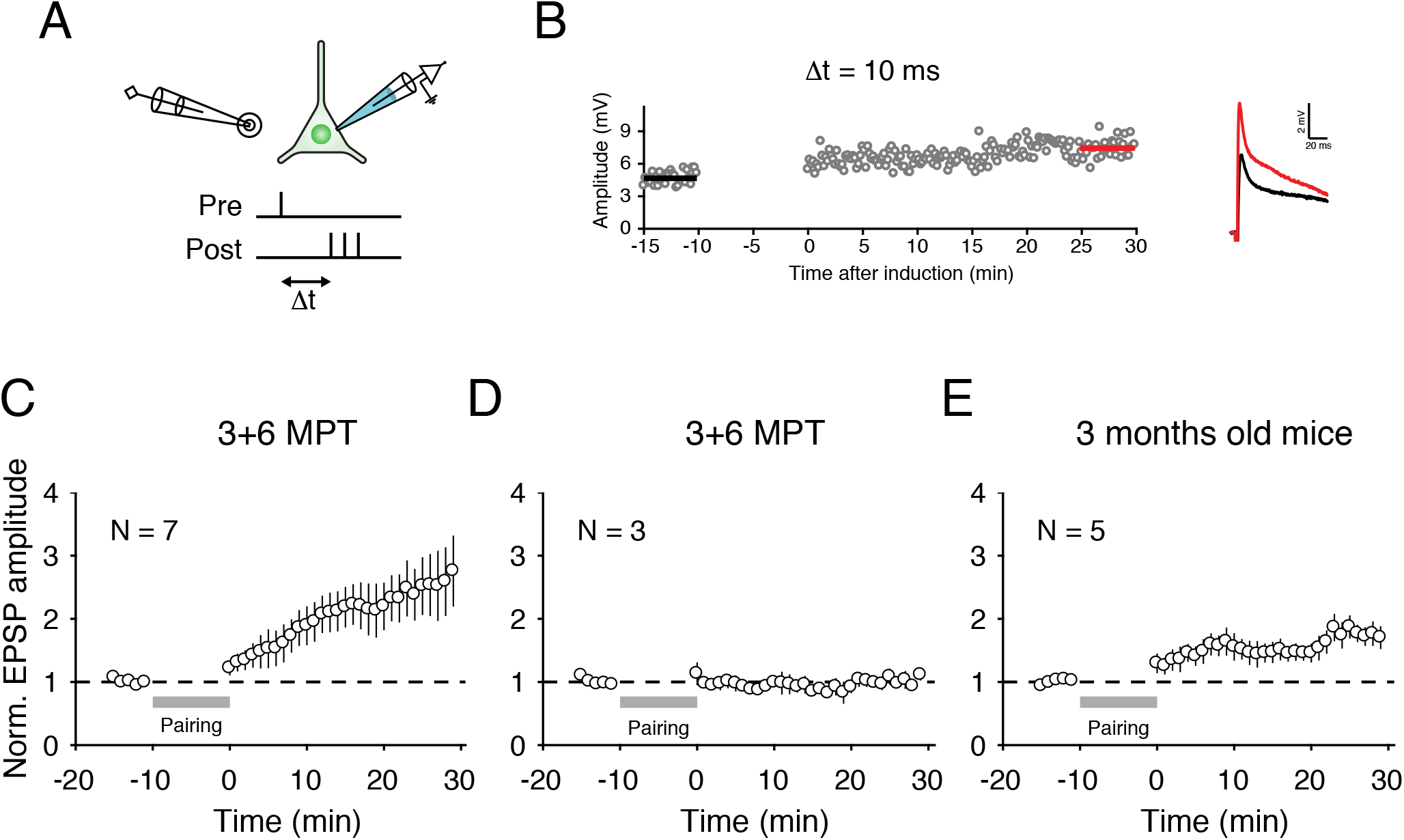
Long-term potentiation of local cortical inputs to hESC-derived neurons in the mouse cortex. (A) Schematic representation of pipette locations and relative timing of extra- and intra-cellular stimulations. (B) Example of one representative cell undergoing potentiation over the 45 minutes of duration of the protocol. (C) EPSP amplitude as a function of time in response to 10 minutes of pre-post pairing for the N=7 human cells that displayed LTP. (D) Same as C, but for the N=3 human cells that displayed a stable EPSP amplitude in response to the pairing protocol. (E) Same as C and D, but for N=5 mouse cells recorded in 3 months-old mice. Markers and error bars indicate mean ± SEM.

### Transplanted human neurons display decorrelated activity and responses to visual stimuli similar to those of host cortical neurons

Synaptic integration within the host cortex may allow xenotransplanted neurons to respond to internal and external events and to encode sensory information. To assess the *in vivo* response features of xenotransplanted cortical neurons, we performed cellular calcium imaging (Figure 6). Using a two-photon microscope and a chronically implanted glass window, we imaged in layer 2/3 of visual cortex the somatic calcium responses of xenotransplanted neurons expressing both the calcium indicator GCaMP6s and nuclear label nls-tdTomato (Figure 6A). We examined during quiet wakefulness spontaneous somatic calcium transients, and visually-evoked activity in response to drifting gratings stimuli (Figure 6B). GCaMP6s expression was driven conditionally 2 weeks prior to imaging by Doxycyclin to prevent any developmental defects, while the constitutive expression of the nuclear label allowed identification of transplanted neurons. We focused on cells located in visual areas of posterior cortex, 100-300 *µ*m below the cortical surface and performed imaging at 5-9 MPT.

**Figure 6.**
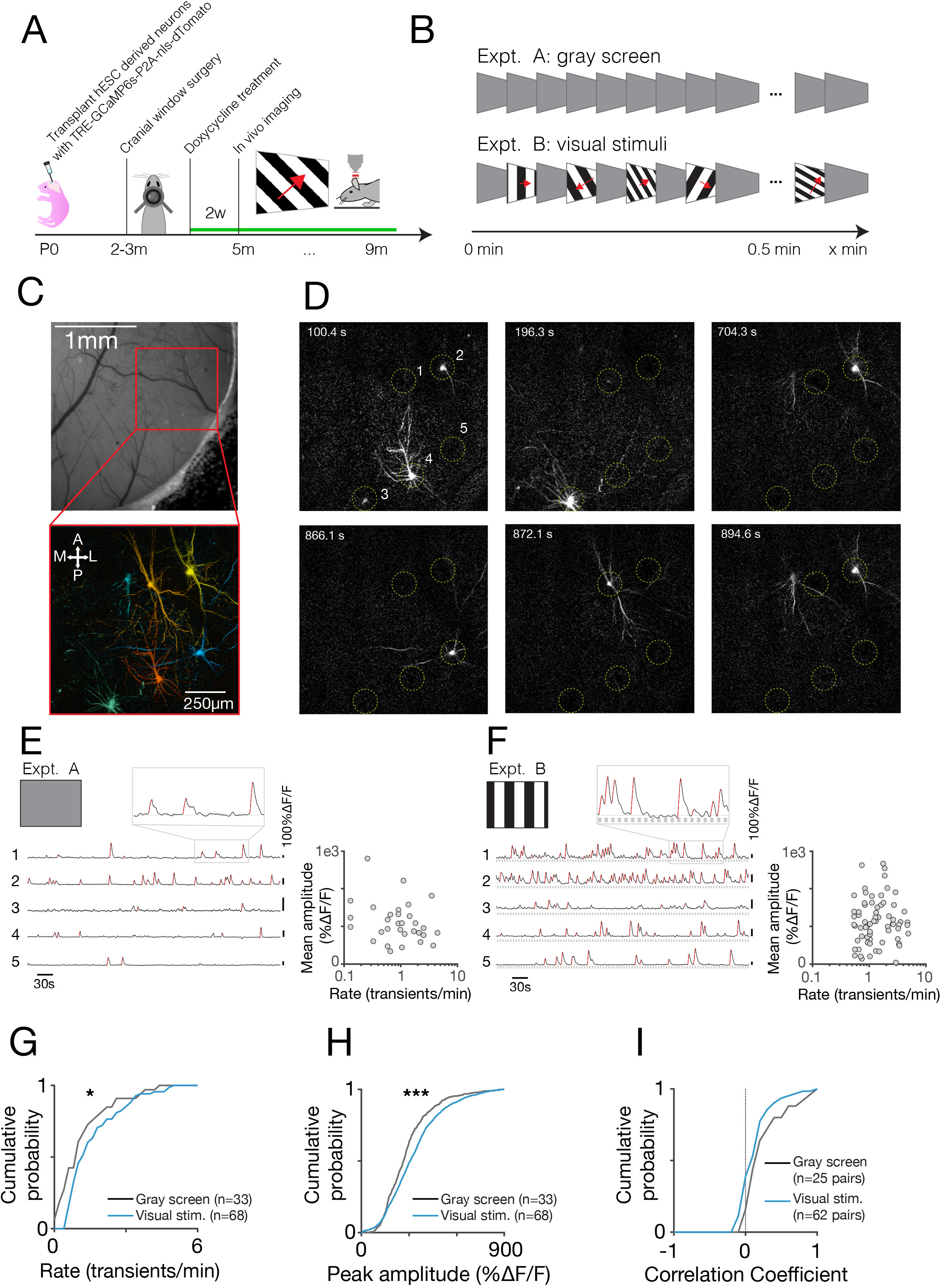
Functional imaging reveals functional responses to visual stimuli. A) Experimental timeline for *in vivo* calcium imaging experiments. Differentiated hESC cells, infected with lenti-TRE-GCaMP6s-P2A-nls-dTomato, are transplanted when pups are born. We perform a cranial window when the animals are 2-3 months old. Two weeks before the experiments start, we induce the expression of GCaMP6s and dTomato using food pellets containing 625 mg/kg doxycycline. (B) Overview of the visual stimulation paradigm. In experiment A, we present a static gray screen to the contralateral eye of head-fixed animals. For experiment B, we present square wave gratings, drifting in 12 direction at different combinations of temporal and spatial frequencies. (C) Example recording results. *Top*: cropped wide-field fluorescence image of the cranial window. Red zoom box shows the location of 2-photon field-of-view shown below. *Bottom*: Projection of hESC neurons recorded using volumetric 2-photon calcium imaging. Neurons are color-coded based on their activity during experiment B. (D) Example data showing the activity of the neurons shown in C) at single frame resolution at different times during experiment B. Yellow dashed circles mark the location of five active neurons. Note how different neurons are active at different moments in time. (E) Spontaneous activity traces for the active neurons shown in C-D. *Left*: Relative fluorescence of single neurons during experiment A (black lines). Positive transients that are used during subsequent quantifications are marked with red lines. The gray zoom box shows a section of the trace at higher magnification. *Right*: Scatter plot showing the quantification of rate and average amplitude for 31 hESC neurons. (F) Activity traces recorded during visual stimulation. *Left*: Relative fluorescence for the same five neurons shown in (E). Epochs of visual stimulation are indicated by gray squares. Note the increased number of transient event marked by red lines. *Right*: Scatter plot showing for 81 neurons that both rate and amplitude are increased when visual stimuli are presented compared to the blank experiment. (G) Comparison of transient rate during gray screen (gray line, N=31 neurons) vs. visual stimulation (blue line, N=81 neurons). Rate of transient increase during visual stimulation. (H) Comparison of transient amplitude gray screen (gray line) vs. visual stimulation (blue line). The amplitude of transients increases during visual stimulation. (I) Comparison of noise (gray line, N=22 pairs) and signal (blue line, N=97 pairs) correlations. Pairwise correlations decrease slightly during visual stimulation.

We observed robust spontaneous and visually-driven activity in more than a quarter of sampled xenotransplanted cells (Figure 6C-F). We measured the activity of 155 transplanted neurons measured from 5 animals (N=54 sessions during visual stimulation, for N=15 sessions also spontaneous activity was recorded). In absence of a visual stimulus (gray screen, Expt. A), about 32% of identified cells showed significant spontaneous somatic calcium transients (>0.5 transients/min, N=25/77 labeled neurons; Supplementary Table). During visual stimulation, a slightly higher fraction, about 35% of cells, showed calcium activity (N=55/155 labeled neurons, Expt. B). Amongst cell showing calcium activity, visual stimulation increased the rates at which calcium transients occurred (Figure 6G; Supplementary Table S1) as well as the peak amplitude of the transients (Figure 6H; Supplementary Table S1). Thus, xenotransplanted neurons not only fire spontaneously but also receive significant inputs from host neurons, which could be from neighboring cortical neurons or from thalamic neurons.

Notably, the activity of xenotransplanted neurons was highly de-correlated (Figure 6D,I, Figure S6C,D). Simultaneously imaged xenotransplanted neurons fired independently, with distinct cells showing calcium transients at distinct time points (Figure 6D). Accordingly, calcium activity time courses in presence and absence of visual stimulation were only weakly correlated (Supplementary Table S1). This suggests xenotransplanted neurons develop to receive highly specific inputs, both from the eyes and other sources.

To examine the specificity of these inputs, we examined tuning of responses for grating direction and orientation (Figure 7). Neurons in mouse visual cortex are characterized by diversely tuned responses to visual stimuli encoding, for example, stimulus orientation and direction (Niell and Stryker, 2008). If human neurons receive inputs from host neurons then they should also show such diversely tuned responses to visual stimuli.

**Figure 7.**
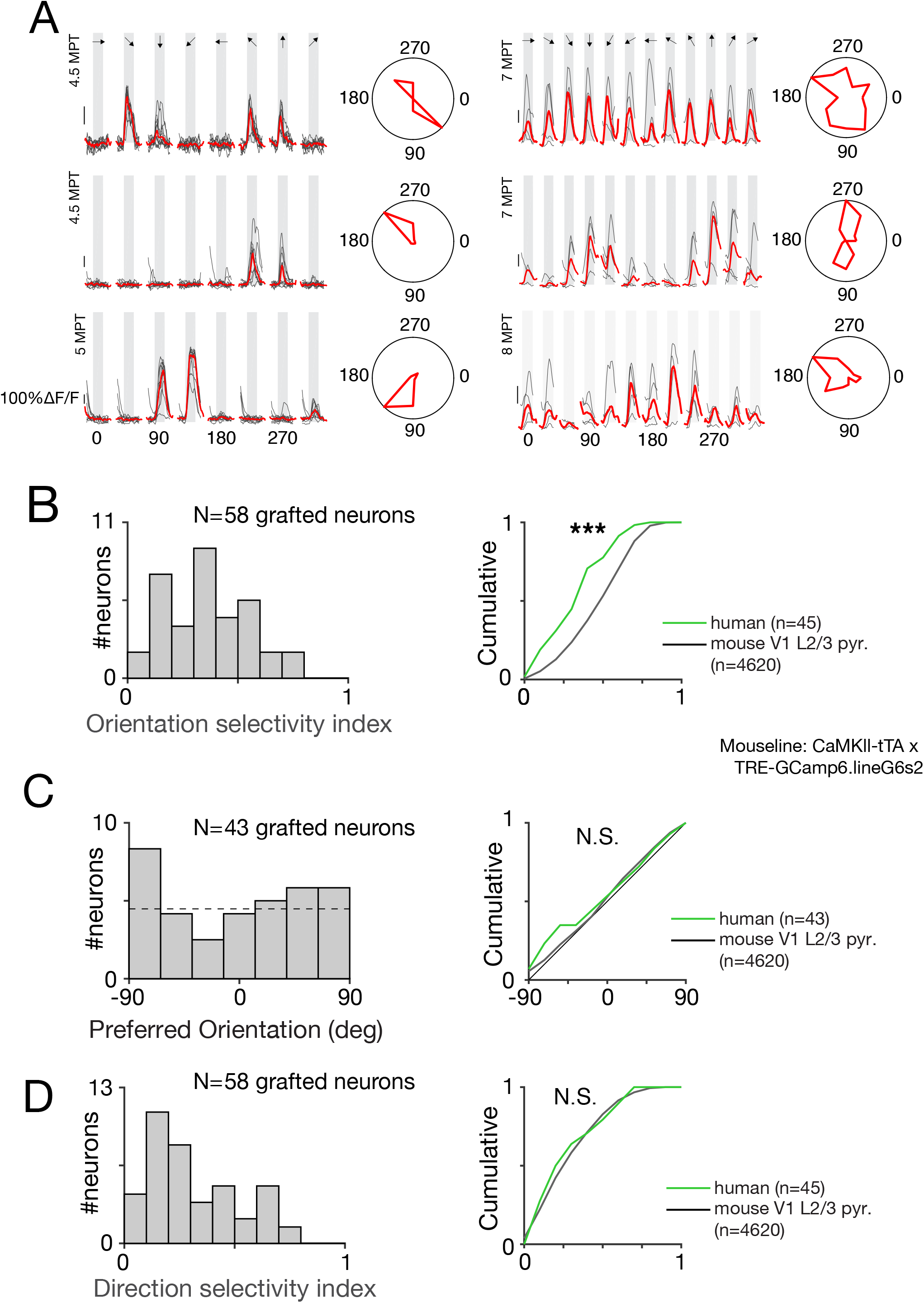
Visual responses are physiologically tuned and similar to those of mouse neurons. A) Example responses to drifting gratings for 6 hESC neurons. Left: neurons recording in Exp. A. Right: neurons recorded in Exp. B (showing best SF-TF combination, see methods). Each sub-panel consists of sorted responses to different orientations; grey bars indicate stimulus presentation. To summarise the tuning profile, the same data are also shown in a polar plot on the right. Some properties of this profile are quantified in panels B-D. (B) Summary of orientation selectivity indices. *Left*: Distribution of orientation selectivity of hESC neurons. *Right*: comparison to responses recorded from mouse V1 L2/3 pyramidal neurons (CaMKII-tTA x TRE-GCaMP6 line G6s2). (C) Summary of preferred orientation. *Left*: Distribution of preferred direction of hESC neurons. *Right*: comparison between responses recorded from hESC (green line) and mouse (gray line) V1 L2/3 pyramidal neurons. (D) Summary of direction selectivity indices. Left: Distribution of direction selectivity of hESC neurons. Right: comparison to responses recorded in from mouse V1 L2/3 pyramidal neurons.

About 25% of xenotransplanted neurons (38/155 labeled neurons) showed clear visual responses (median ΔF/F_0_ response > 3 SD over baseline for >1 second). Remarkably, responses were diversely tuned for different directions of motion and spatial orientations (Figure 7A). To quantify these responses properties, we computed the stimulus direction and orientation that elicited the strongest response (preferred direction and orientation), and indices quantifying the selectivity of responses for these stimulus attributes (direction and orientation selectivity indices: DSI and OSI respectively). We compared these data to those that we obtained from mouse layer 2/3 pyramidal neurons (N=4260, mouse line: CaMKII-tTA x TRE-GCaMP6 line G6s2).

Consistent with the weak activity correlations observed between cell pairs, xenotransplanted neurons showed diversely tuned responses (median OSI=0.36 and DSI=0.25; Figure 7B-D, left) with tuning properties resembling those of mouse neurons (Figure 7B-D, right, Figure S5A-C). Selectivity for orientation and direction was pronounced, with 60% and 78% of visually responsive cells showing tuned responses for stimulus orientation and direction (orientation and direction selectivity indices >0.2; Figure 7B&D), similar to that observed in mouse neurons (Supplementary Table S1). Compared to mouse neurons, visual responsive human neurons did show weaker selectivity for orientation but not for motion direction (Supplementary Table S1). This could reflect the maturing nature of the underlying circuitry, which in transplanted mouse neurons shows a sharpening of tuning over time (Falkner et al. 2016). These findings indicate that xenotransplanted human neurons successfully integrate to inherit functions of the host cortical network, including tuned responses to sensory stimuli.

## Discussion

The mechanisms underlying the development of human neurons and synapses have major implications for human brain evolution, function, and disease. Here we take advantage of a novel mouse-human chimeric model to recapitulate key features of human cortical pyramidal neuron development in vivo, with unprecedented cell resolution and functional depth.

While in vitro PSC-based models, whether with adherent cultures or organoids, hold great promises to study experimentally human neural development (Astick and Vanderhaeghen, 2018; Di Lullo and Kriegstein, 2017; Lancaster and Knoblich, 2014), it has remained challenging to apply them to the study of neuronal maturation and function, mostly because of the challenges associated with long-term culture needed to achieve sufficient levels of survival and maturation. Models of xenotransplantation of human neurons in the mouse cortex have been described that constitute an attractive alternative for modelling of human neuron development and disease (Espuny-Camacho et al., 2017; Espuny-camacho et al., 2013; Espuny-Camacho et al., 2018; Mansour et al., 2018; Real et al., 2018). However, while these models revealed the ability of the neurons to integrate morphologically and send robust and specific axonal projections, their level of functional integration within cortical circuits has been limited by the fact that in these models most transplanted cells aggregate in clusters, thus in a microenvironment quite distinct from the mouse brain.

In contrast to these models, here we find that when human neurons can integrate as single cells within mouse cortical circuits, they display strong connectivity with the host neurons, as shown from rabies virus tracing, combined electrophysiology and morphology, as well as in vivo structural and functional imaging.

This approach recapitulates in a robust fashion several key milestones of human neuronal development that have not been reported so far, whether in other models of xenotransplantation or in vitro models such as monoadherent cultures or neural organoids (Di Lullo and Kriegstein, 2017; Suzuki and Vanderhaeghen, 2015). Specifically the transplanted neurons displayed (1) coordinated neuronal morphogenesis and physiological maturation at the single cell level, (2) robust dendritic spine dynamics followed by stabilization and synaptogenesis (3) functional synaptic plasticity, (4) physiological responses to sensory stimulation that are similar to those of native cortical neurons.

We and others previously reported a strikingly prolonged time-line of morphological development for transplanted human cortical pyramidal neurons and interneurons, that we proposed to be in line with human cortical neoteny (Anderson and Vanderhaeghen, 2014; Espuny-camacho et al., 2013; Maroof et al., 2013; Nicholas et al., 2013). However, the exact temporal pattern of synaptic maturation and functional integration of xenotransplanted human cortical neurons had not been characterized systematically. Moreover, it had remained unclear whether the prolonged synaptic maturation of human neurons xenotransplanted in the mouse cortex is genuinely intrinsic to these cells, or rather a collective property of transplanted cells remaining mostly isolated from the rest of the brain. Here we find that xenotransplanted neurons integrated as single cells within the mouse cortex developed morphologically and functionally in a coordinated fashion, like in vivo, and this developmental program follows a months-long timeline that is remarkably similar to the one reported in the human developing brain (Defelipe, 2011; Sousa et al., 2017). Most strikingly, the human neurons display high levels of dendritic spine structural dynamics typical of juvenile neurons even 6 months following transplantation. In contrast, dendritic spines of mouse neocortical neurons display high dynamics and turnover only for the first few weeks postnatally, after which they stabilize during adulthood, with much more limited turnover (Hofer et al., 2009; Holtmaat and Svoboda, 2009). Our observation that human neuron dendritic spines display highly dynamic patterns in the adult mouse host cortex thus provides strong evidence that the prolonged development of human cortical neurons is not likely to be an experimental artifact, but rather a robust biological feature of human neurons, with a strong intrinsic component. This intrinsic retention of juvenile properties of human cortical neurons may constitute the cellular basis of the long-known brain neoteny that is thought to have played a major role in human evolution. Importantly, our data now provide a framework for the design of mechanistic studies aimed at deciphering the cellular and molecular mechanisms underlying the striking neotenic nature of human cortical neurons.

Notably, despite their prolonged timeline of developmental dynamics, the transplanted neurons are not stuck in this immature state. Instead they gradually display increased spine density and decreased spine turnover, as would be expected during normal development, and finally reach levels of spine stability and density that are close to those of mouse adult neurons. This contrasts with findings recently reported on PSC-derived human cortical neurons transplanted as bulk cell populations, that displayed rapid dendritic spine turnover throughout the experiment (spine half-life less then 4 days vs. 2 weeks at 3-4 months), and reached very limited levels of synaptic connectivity with the host brain (Real et al., 2018). These differences illustrate the important influence of the transplantation paradigm, in particular the ability of the transplanted neurons to integrate as single cells, on the levels of maturation and connectivity that can be achieved by the transplanted neurons. The xenotransplanted neurons also reach much more advanced stages of spine synapse maturation than reported so far with in vitro models of human corticogenesis, whether using adherent cultures or organoids (Astick and Vanderhaeghen, 2018; Di Lullo and Kriegstein, 2017; Lancaster and Knoblich, 2014), which suggests that the host brain provides not only a permissive environment for long term-maturation, but also instructive cues for complex events such as spine morphogenesis and synaptogenesis.

Importantly, the high level of connectivity displayed by the transplanted neurons enabled to detect robust functional synaptic plasticity, i.e., long term potentiation, which was to our knowledge never reported before for human PSC-derived neurons. As long term potentiation is a major read-out of synaptic function within a whole circuit, this provides unique opportunity to exploit this in stem cell-based modelling of human neuronal plasticity, especially in the context of brain disease modelling. Indeed, synaptic function and plasticity are cardinal features of pyramidal neurons that are thought to be altered in many neurodevelopmental and degenerative diseases (Cruz-Martin et al., 2010; Ebert and Greenberg, 2013; Murmu et al., 2013; Spires-Jones and Hyman, 2014; Zoghbi and Bear, 2012).

On the other hand it is important to note that, while the xenotransplanted human neurons described here have achieved unprecedented maturation, their morphological and functional properties still correspond to a less mature stage than adult human neurons. For instance, morphologically the transplanted human neurons display smaller cell body and dendritic arbours than adult human cortical neurons, and functionally only a fraction of the neurons display visual reponses among which many show sligthly weaker orientation tuning, in line with the fact that human cortical neurons are thought to take up to several years to reach full maturity. Similarly, the dendritic spine dynamics analyses suggest that the neurons are still undergoing synaptic growth and have not yet reached the stage of dendritic spine/synapse pruning, which is supposed to last for more than decade in some areas of the human neocortex (Petanjek et al., 2011). It will be interesting to test in the future whether transplanted human neurons allowed to develop for longer periods can eventually acquire functional and morphological properties of mature human neurons such as larger dendritic arbors and spine density (Benavides-Piccione et al., 2002; Petanjek et al., 2011; Rakic et al., 1986), higher capacity of transmitting information (Beaulieu-Laroche et al., 2018; Eyal et al., 2016; Testa-Silva et al., 2014), and an increased electrical compartmentalization (Beaulieu-Laroche et al., 2018).

Most importantly, the model provides a first characterization of the function of human neurons in an intact circuit in vivo. Indeed we report that up to 25% of the xenotransplanted neurons display functional responses to visual sensory stimulation, similar to those of host neurons. Remarkably, these neurons fire in a decorrelated fashion and display visually tuned responses that resemble the responses of mouse cortical neurons (Andermann et al., 2011). Similar results for visual tuning are reported for mouse neurons transplanted into lesioned adult mouse cortex 3-4 months after transplantation, which show OSI and DSI values that resemble those of native mouse neurons (Falkner et al., 2016). This shows that xenotransplanted human neurons can integrate in the adult mouse brain and gain information processing function that will be useful in studies of cortical development and repair.

These data also constitute a striking example of the capacity of single neurons to adapt to the dominant influence of the circuit in which they are integrated. It will be fascinating in future studies to decipher the mechanisms involved, focusing on the acquisition and tuning of sensory responses, and determine whether and how they compare with those of host neurons that normally occur very early on in the mouse (Mizuno et al., 2018). In this respect it will be also interesting to explore in the future any difference that may be related to the distinct anatomical organization and development of tuning specificity between the mouse and primates (Van den Bergh et al., 2010).

Another surprising aspect of our *in vivo* imaging dataset is that genuine functional integration of the transplanted neurons was observed mainly at an advanced adult stage (>6MPT), implying that juvenile neurons could engage functionally into a mature cortical circuit in relatively old animals, and yet display response highly similar to those of the host neurons. This has potentially important implications in the perspective of brain repair by cell therapy strategies (Tabar and Studer, 2014). Several models of transplantation of human PSC-derived cortical neurons into lesioned adult cortex have been shown to be promising in this context (Tornero et al., 2017; Tornero et al., 2013). It will be interesting to test the transplantation method described here in such experimental models of cortical lesions, to assess whether and how transplanted neurons can integrate into damaged cortical circuits, up to the functional level shown here following neonatal transplantation, as an important step towards the restoration of damaged cortical circuits.

## Acknowledgments

The authors thank members of the Vanderhaeghen lab for helpful discussions and advice and J.-M. Vanderwinden of the ULB Light Microscopy Facility for his support with imaging. A subset of the images were acquired on a Zeiss LSM 880 – Airyscan system (Cell and Tissue Imaging Cluster, CIC), supported by Hercules AKUL/15/37_GOH1816N and FWO G.0929.15 to Pieter Vanden Berghe, University of Leuven. This work was funded by the Vlaams Instituut voor Biotechnologie (VIB), Neuro-Electronics Research Flanders, the European Research Council (ERC Adv Grant GENDEVOCORTEX) (to P.V.), KU Leuven Research Council Grant C14/16/048 (to V.B.), the EOS Program (to P.V.), the Belgian FWO and FRS/FNRS (to P.V.), the WELBIO Program of the Walloon Region (to P.V.), the AXA Research Fund (to P.V.), the Fondation ULB (to P.V.). R.I. was supported by a postdoctoral Fellowship of the FRS/FNRS, and B.L.P. by a postdoctoral Fellowship of the FWO.

## Author Contributions

Conceptualization and Methodology, D.L., B.V., R.I., V.B., and P.V.; Investigation, D.L., B.V., A.R., B.A.D., L.B., B.L.P., A.B., L.D., D.G., R.I., V.B., and P.V.; Formal Analysis, D.L., B.V., R.I., V.B., and P.V.; Key reagents, K.C. Writing - Original Draft, D.L., B.V., V.B., and P.V.; Writing - Review & Editing, D.L., B.V., A.R., B.D., D.L., D.G., R.I., V.B., and P.V.; Funding acquisition, P.V.; Resources, V.B. and P.V.; Supervision, V.B. and P.V.

## STAR Methods

### KEY RESOURCES TABLE

Separate file

### CONTACT FOR REAGENT AND RESOURCE SHARING

Further information and requests for resources and reagents should be directed to and will be fulfilled by the lead contact.

### EXPERIMENTAL MODEL AND SUBJECT DETAILS

#### Animals

All mouse experiments were performed with the approval of the KULeuven and ULB Committees for animal welfare. Animals were housed under standard conditions (12 h light:12 h dark cycles) with food and water *ad libitum*. Data for this study are derived from a total of 65 mice of both sexes.

#### Human ESC differentiation into cortical cells

Human ESC H9 (Thomson et al., 1998) and H9-GFP (Espuny-camacho et al., 2013) cells were maintained on irradiated mouse embryonic fibroblasts (MEF) in the ES medium until the start of cortical differentiation. Cortical differentiation from human ESC was performed as described previously (Espuny-camacho et al., 2013) with some modifications (Suzuki et al., 2018). On day −2, ESCs were dissociated using Stem-Pro Accutase (Thermo Fisher Scientific, Cat#A1110501) and plated on matrigel- (hES qualified matrigel BD, Cat#354277) coated dishes at low confluency (5,000–10,000 cells/cm2) in MEF-conditioned hES medium supplemented with 10 *µ*M ROCK inhibitor (Y-27632; Merck, Cat#688000). On day 0 of the differentiation, the medium was changed to DDM (Gaspard et al., 2008), supplemented with B27 devoid of Vitamin A (Thermo Fisher Scientific, Cat#12587010) and 100 ng/ml Noggin (R&D systems, Cat#1967-NG), and the medium was changed every 2 days until day 6. From day 6, the medium was changed every day until day 16. After day 16 of differentiation, the medium was changed to DDM, supplemented with B27 (DDM/B27), and changed every day. At day 25, the progenitors were dissociated using Accutase and cryopreserved in mFreSR (StemCell Technologies, Cat#05855).

For comparison cortical neurons derived with another protocol, cortical differentiation was performed as in (Shi et al., 2012).

Differentiated cortical cells were validated for neuronal and cortical markers by immunostaining using antibodies for TUBB3 (BioLegend, Cat#MMS-435P), TBR1 (Abcam, Cat#ab183032), CTIP2 (Abcam, Cat#ab18465), and FOXG1 (Takara, Cat#M227).

#### Viral constructs

HEK293T cells were transfected by packaging plasmids, psPAX2 (Addgene Cat#12260) and pMD2.G (Addgene Cat#12259), and a plasmid of gene of interest in lentiviral backbone (pLenti-hSynapsin I promoter-EmGFP-WPRE, pLenti-CAG-Venus-WPRE, pLenti-hSynapsin I promoter-Venus-WPRE, pLenti-hSynapsin I promoter-TVA-mCherry–P2A–N2c(G)-WPRE, pLenti-TRE-GCaMP6s-P2A-nls-dTomato-WPRE and FUW-M2rtTA(Addgene Cat#20342)). 3 days after transfection, culture medium was collected and viral particles were enriched by filter device (Amicon Ultra-15 Centrifuge Filters, Merck, Cat#UFC910008). Titer check was performed on HEK293T cell culture for every batch of lentiviral preparation.

#### Neonatal transplantation

Human cortical cells that were frozen at day 25, were thawed and plated on matrigel-coated plates using DDM/B27 and Neurobasal supplemented with B27 (DDM/B27+Nb/B27) medium. Seven days after plating, cells were dissociated using Accutase and plated on new matrigel-coated plates at high confluency (100,000–600,000 cells/cm2) with or without lentiviral vector. The following day, medium was changed to DDM/B27+Nb/B27 medium. At 14-16 days after thawing, cells were treated with 10 *µ*M DAPT (Abcam, Cat#ab120633) for 24 hours. The following day, cells were treated with 5 *µ*M Cytarabine (Merck, Cat#C3350000) for 24 hours. At day 17-19 days after thawing, cells were dissociated using Accutase and suspended in the injection solution containing 20 mM EGTA (Merck, Cat#03777) and 0.1% Fast Green (Merck, Cat#210-M) in PBS at 40,000–200,000 cells/*µ*l. Approximately 1-2 *µ*l of cells’ suspension was injected into the ventricles of each hemisphere of neonatal (postnatal day 0 or 1) immunodeficient mice (NOD/SCID or Rag2^-/-^) using glass capillaries pulled on a horizontal puller (Sutter P-97).

#### Electrophysiology

Whole-cell patch-clamp recordings were performed intracellularly on grafted neurons in acute brain slices from mice starting at 1 and up to 11 MPT, as previously described (Espuny-Camacho et al., 2013). Briefly, transplanted mice were lightly anesthetized with isofluorane and decapitated. Alternatively, animals older than 3 months were anesthetized intraperitoneally with Nembutal and transcardially perfused with ∼20 ml of ice-cold NMDG-based slicing solution (Jiang et al., 2015; Ting et al., 2014). Brains were rapidly extracted and placed in ice-cold NMDG-based slicing solution containing (in mM): 93 *N*-Methyl-D-glucamine, 2.5 KCl, 1.2 NaH_2_PO_4_, 0.5 CaCl_2_, 10 MgSO_4_, 30 NaHCO_3_, 5 Na-ascorbate, 3 Na-pyruvate, 2 Thiourea, 20 HEPES and 25 D-glucose (pH adjusted to 7.35 with 10 N HCl, gassed with 95% O_2_/5% CO_2_). The cerebellum and hindbrain were trimmed and the remaining block of tissue was glued onto the slicing platform of a Leica VT1200. 250-300 *µ*m thick coronal slices were cut in ice-cold NMDG-based slicing solution and subsequently incubated for ∼10 minutes in the same solution at 36°C. Slices were then stored for at least 45 minutes at room temperature in standard aCSF, containing (in mM): 125 NaCl, 2.5 KCl, 1.25 NaH_2_PO_4_, 2 CaCl_2_, 1 MgCl_2_, 30 NaHCO_3_ and 25 D-glucose (gassed with 95% O_2_/5% CO_2_). For recording one slice at a time was transferred to a chamber and continuously perfused with aCSF at ∼1 ml/min. Transplanted cells were identified by their EGFP fluorescence and visualized using an upright microscope equipped with infrared differential interference contrast (Carl Zeiss NV, Belgium). Whole-cell patch-clamp recordings and bipolar extracellular stimulation were performed and analysed as described in more detail in the Supplemental Experimental Procedures.

#### Rabies virus-based monosynaptic tracing

For rabies-based monosynaptic tracing, human cortical cells frozen at day 25 were thawed and infected in vitro with lenti-hSynI-TVA-mCherry-P2A-N2c(G) at day 32 (transfection efficiency ∼50%). Cells were subsequently grafted at day 41 into the P0 mouse cortex following the previously described procedure. At approximately 3 MPT, n=2 NOD/SCID mice were anaesthetized with isoflurane (2.5%–3% induction, 1%–1.25% maintenance) and a small craniotomy approximately 2 mm in diameter targeting barrel cortex was performed 0.7 mm posterior and 3.5 mm lateral from Bregma. Approximately 200 nl of rabies virus (N2c ΔG-eGFP(EnvA)) were injected using a bevelled glass capillary mounted on a Nanoject II injector (Drummond Scientific) at a depth of 0.45 mm from the brain surface at a rate of 18.4 nl per injection with ∼30 sec between injections. After 2 weeks, animals were perfused with freshly prepared 4% paraformaldehyde (Invitrogen) and subsequent histology was performed as described in the Immunofluorescence section.

#### Surgical procedures

Standard craniotomy surgeries were performed to gain optical access to the visual cortex through a set of cover glasses (Goldey et al., 2014). Rag2KO mice aging between 2 and 6 months were anaesthetised (isoflurane 2.5%–3% induction, 1%–1.25% or a mix of ketamine and xylazine 100mg/kg and 10 mg/kg respectively). A custom-made titanium head plate was mounted to the skull, and a craniotomy over the visual cortex was made for calcium imaging. The cranial window was covered by a 4 or 5 mm cover glass. Buprenex and Cefazolin were administered postoperatively (2 mg/kg and 5 mg/kg respectively) when the animal recovered from anaesthesia after surgery.

#### Widefield Calcium Imaging

Widefield fluorescent images were acquired through a 2x objective (NA = 0.055, Edmund Optics). Illumination was from a blue LED (479nm, ThorLabs), the green fluorescence was collected with an EMCCD camera (EM-C2, QImaging) via a bandpass filter (510/84 nm filter, Semrock). The image acquisition was controlled with customized software written in Python.

#### Two-photon Calcium Imaging

A customized two-photon microscopy (Neurolabware) was used. GCaMP6s were excited at 920 nm wavelength with a Ti:Sapphire excitation laser (MaiTai eHP DeepSee, Spectra-Physics). The emitted photons were split by a dichroic beamsplitter (centered at 562 nm) and collected with a photomultiplier tube (PMT, Hamamatsu) through a bandpass filter (510±42 nm, Semrock) for the green fluorescence of GFP or GCaMP6s and a bandpass filter (607±35 nm, Semrock) for the red fluorescence of nls-dTomato.

#### In vivo structural imaging

Animals were aneasthetised using a mix of ketamine (100 mg/kg) and xylazine (10mg/kg) at 1% ml/g of their body weight. They were placed on a sterilized plexiglass platform and protective eye ointment was applied to both eyes.

The high NA objective (25x Olympus, 1.05 NA) was aligned to be orthogonal to the surface of the top coverslip. All imaging was performed by moving the motorised stages of the microscope along a virtual axis parallel to the optical axis of the objective. We acquired high resolution anatomical stacks from anaesthetised mice. Typical stacks consisted of 200-300 optical section spaced 1 *µ*m apart. The imaged area spanned 75×120 *µ*m. To reduce effects of shot noise, we averaged over 50 frames collected per section and applied a 3d median filter (Fiji, kernel size=[1.5, 1.5, 4]) over the entire stack. Using these preprocessed stack, we traced branches of interest using a custom matlab GUI. We then extracted 3D sub-volumes (10 x 10 x X *µ*m, where X is the length of the branch, ranging 50-120 *µ*m) rotated in all 3 dimension to closely fit each branch. These sub-volumes were used for subsequent analysis.

#### In vivo functional imaging

Two-photon images (702×796 pixels per frame) were collected at 20 Hz with a 16x objective (Nikon 0.80 NA). Volume imaging was accomplished by using an electro-tunable lens (EL-10-30-TC, Optotune) to move the focal plane using a sawtooth pattern in 4 or 5 steps (50 *µ*m separation). We simultaneously recorded neuronal activities in large volumes (1 x 1.5 x 0.20 mm^3^) of layer 2/3 visual cortex. During imaging, mice were head-clamped on a sterile plexiglass platform while consciously viewing the visual stimuli on the display. Eye movements were monitors using a camera and infrared illumination (720–900 nm bandpass filters, Edmund).

Visual stimuli were displayed on a gamma-corrected LCD display (22’’, Samsung 2233RZ). The screen was oriented parallel to the eye and placed 18 cm from the animal (covering 80 degree in elevation by 105 degree in azimuth). Spherical correction was applied to the stimuli to define eccentricity in spherical coordinates. Visual stimuli consisted of drifting square wave gratings in 6 combinations of 3 spatial frequencies (SF=0.04, 0.08, 0.16 cycles per degree) and 2 temporal frequencies (TF=1, 4 Hz) in 12 directions covering 360 degrees. In an early version of the experiment, gratings (SF=0.05 cpd and TF=4 Hz) drifting in 8 directions were presented. We collected a control dataset from the CaMKll-tTA x TRE-GCamp6 line G6s2 transgenic mouse line (Wekselblatt et al., 2016).

#### Immunofluorescence

Mice were fixed by transcardiac perfusion with freshly-prepared 4% paraformaldehyde in PBS (Invitrogen). Brains were dissected, and 100 *µ*m sections were prepared using a Leica VT1000S vibrosector. Slices were transferred into PBS with 0.5 *µ*g/mL sodium azide (Sigma), then blocked with PBS supplemented with 3% horse serum (Invitrogen) and 0.3% Triton X-100 (Sigma) during 1 h, and incubated overnight at 4°C with the primary antibodies. After three washes with PBS/0.1% Triton X-100, slices were incubated in PBS for 1 h at room temperature and incubated 2 h at room temperature with the appropriated secondary antibodies. Sections were again washed three times with PBS/0.1% Triton X-100, stained with Hoechst (bisBenzimide H 33258, Sigma) or DAPI (Sigma) for 60 min and washed twice in PBS. The sections were next mounted on a Superfrost slide (Thermo Scientific) and dried using a brush before adding Glycergel mounting medium (Dako). Imaging was performed using either a Zeiss LSM780 or LSM880 confocal microscope controlled by the Zen Black software (Zeiss).

### QUANTIFICATION AND STATISTICAL ANALYSIS

#### Electrophysiology

All electrophysiological recordings were analyzed using MATLAB (The Mathworks, Natick, MA). Raw voltage traces were filtered at 5 kHz and passive membrane and AP properties were extracted and saved in a database for further analysis and statistics.

#### Spine imaging

Segmented branches for adjacent time points were loaded into a custom GUI and all protrusions where marked as spines. During this process we took the 3d structure of the spine into account by scrolling through the volume. This allowed us to distinguish between actual spines and processing passing above and below the branch of interest. We then correlated spines in both time points using their location relative to branch landmarks. Spines that were present in both time points were marked as *conserved*. Spine present only in the first time point were marked as *lost* and, conversely, spines that were found only in the second time point were marked *gained*. Spines present on a non-overlapping section of the branch were marked, but only included for the spine density quantification. The spine *density* was calculated by counting the number of actual spines on each segment and dividing this count by the length of the branch. This length was obtained by summing the distances between adjacent markers used to trace the branch. Spine *turn-over* was calculated by dividing the sum of gained and lost spines by the sum of spines present at both time points. Spine density for gained and lost spines was calculated by counting gained and lost spines and dividing by branch length. To obtain the survival function, we started by taking all spines present in the first time point (*reference*) and then checking for how many subsequent time points each spine of the set was conserved. Survival fraction for individual time point *t* was then calculated by dividing the total number of surviving spines at time *t* by the total number of spines in the *reference* set.

#### Calcium imaging

Two-photon movies for all experiments collected during 1 session were motion registered to a common reference image. This image was constructed by registering and then averaging 1200 frames from the centre of the session. We manually segmented regions-of-interest (ROIs) using information from both red and green channels. We then extracted cellular time courses for each ROI by averaging all pixels inside each ROI, removing the neuropil signal and correcting for slow baseline drift. We then calculated ΔF/F_o_ traces by subtracting the baseline fluorescence (F_o_) corrected time course and dividing by F_o_.

To quantify activity of neurons, we used an differential algorithm that detected epochs were the calcium trace increased by more than 2*STD/second, where STD is the standard deviation of the distribution constructed by combining the negative part of the trace with its sign-inverted counterpart. Neurons were considered *active* if their calcium trace exhibited more than 0.5 transients per minute. Neurons were deemed *visually responsive* if, for at least one condition, the median response exceeded 3x STD, where STD is the standard deviation of the blank epochs preceding each stimulus epoch for this condition.

For orientation tuning experiment we determined the *orientation selectivity index* (OSI) and *direction selectivity index* (DSI) based on circular variance as proposed by Mazurek 2014 (Mazurek et al., 2014). Neurons with an OSI/DSI > 0.2 were deemed *tuned* for orientation/direction of the drifting grating. For tuned neurons, we calculate the *preferred orientation* as the vector means responses to individual directions.

#### Morphology reconstructions and analysis

At the end of each electrophysiological recording, slices containing a single biocytin-filled cell were placed in freshly-prepared 4% paraformaldehyde and fixed overnight. Slices were stained as described previously, using as primary antibodies chicken anti-GFP (Abcam, 1:2000) and occasionally mouse anti-human nucleus (Merck, 1:500) to confirm the human identity of the recorded cells. Streptavidin conjugate NL557 (R&D Systems, 1:2000) was added to the secondary antibodies, and left to incubate for at least 2 hours at room temperature. Full morphologies were imaged on a Zeiss LSM780 or LSM880 confocal microscope, using 40x (NA 1.3) or 63x (NA 1.4) oil immersion objectives, at a horizontal resolution of 0.088 – 0.138 *µ*m/pixel and a vertical resolution of 0.4 – 0.5 *µ*m/pixel. Zeiss proprietary files were converted to stacks of TIFF files, which were then automatically stitched. The morphologies were reconstructed using the semi-automated procedure provided by ShuTu (Jin et al., 2019), followed by manual curation. Digital representations of cellular morphologies were stored in SWC files and analyzed using the TREES toolbox (Cuntz et al., 2010). The Cell Counter toolbox in FIJI (Schindelin et al., 2012) was used to manually annotate spine positions in 3 dimensions.

#### Statistical analysis

Results are shown as mean±standard error (S.E.M.) of at least three biologically independent experiments. Student’s unpaired *t*-test was used for two group comparisons. Analyses of multiple groups were performed by a one-way or two-way analysis of variance (ANOVA) - as indicated in figure legends - followed by post hoc multiple comparisons using Tukey’s test.

### DATA AND SOFTWARE AVAILABILITY

Analysis scripts are available from the lead author upon request.

## Supplemental Information

**Figure S1.**
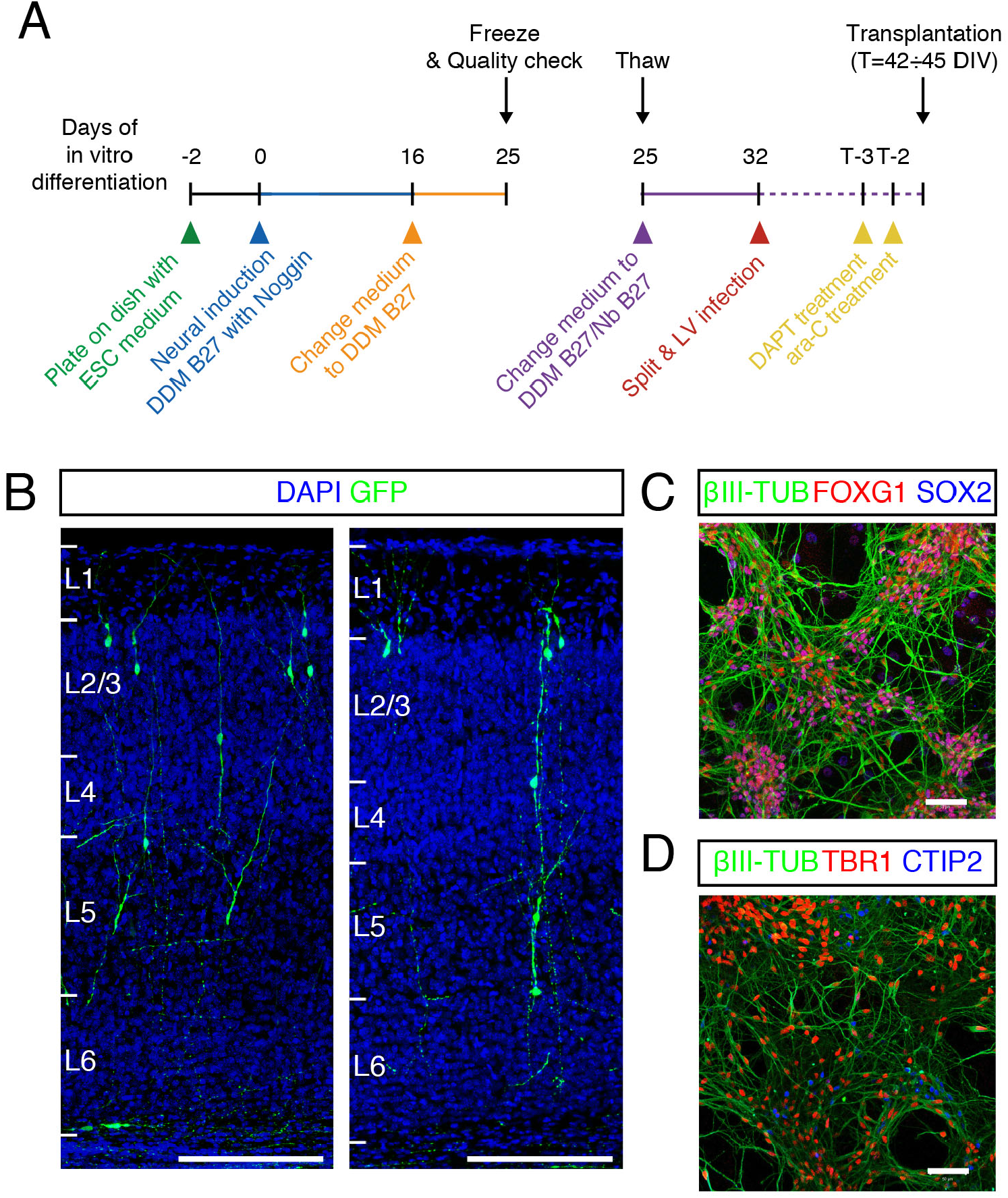
Differentiation protocol. (A) Detailed timeline of the differentiation protocol used to obtain human cortical pyramidal neurons for transplantation in the mouse cortex. (B) Examples of ESC-derived cortical neurons integrated in the mouse cortex throughout the 6 cortical layers. Scale bar, 200 *µ*m. (C,D) In vitro validation of the quality of the differentiated cells: at DIV32, most neuronal cells express markers of cortical identity, including FoxG1, Tbr1, CTIP2. Scale bar, 50 *µ*m.

**Figure S2.**
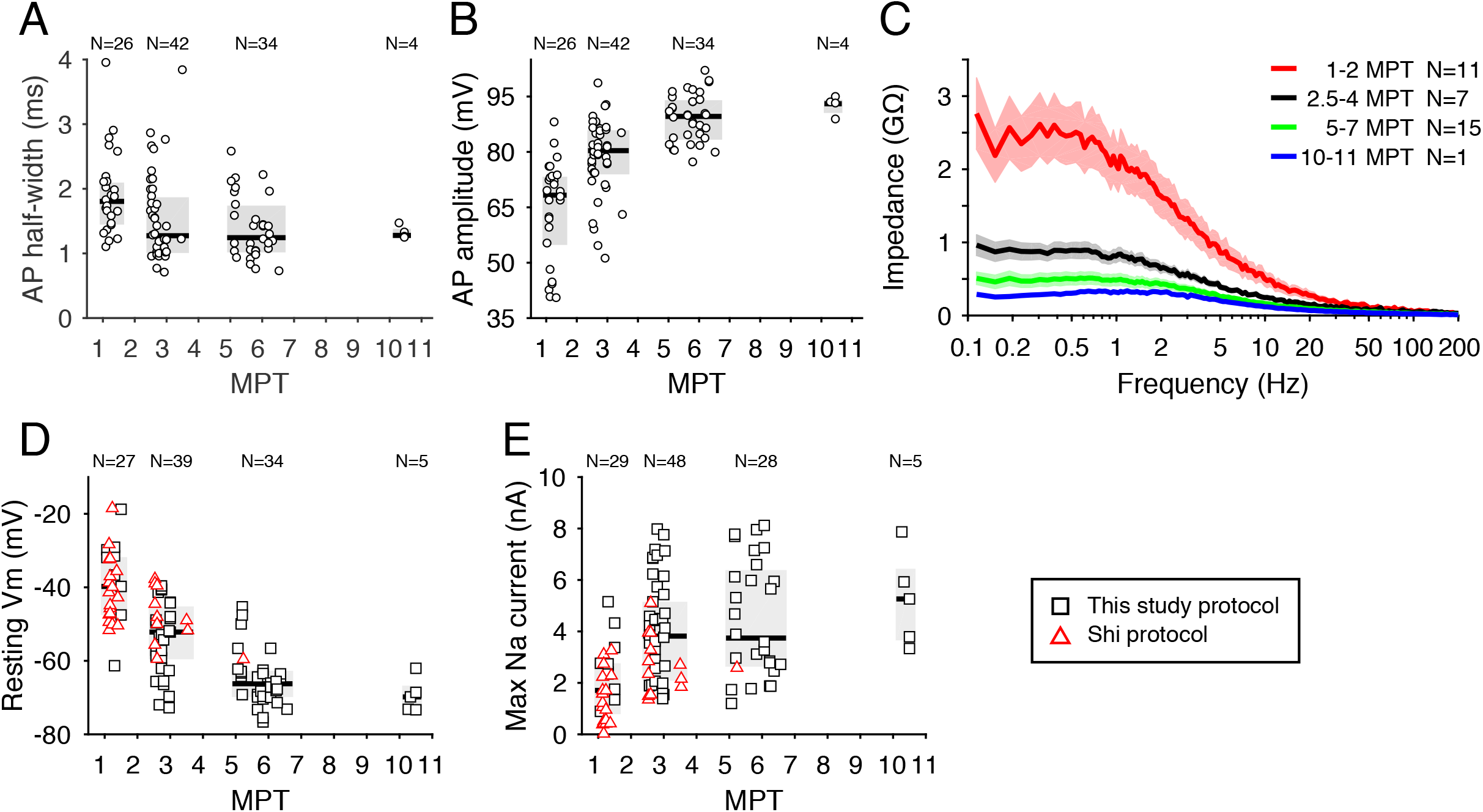
Electrophysiological properties of hESC-derived transplanted cortical neurons. (A,B) Development of action potential half-width and amplitude over the course of 11 MPT. Values attained at the end of this developmental period are comparable to those observed in human ex vivo samples. (C) Membrane impedance as a function of stimulating frequency at 4 consecutive developmental stages. Notice how the overall impedance decreases with time, in line with the reduction of input resistance observed in the same cells. (D,E) Development of resting membrane potential and maximum recorded sodium currents as a function of time, for two distinct differentiation protocols. Notice how the choice of differentiation protocol does not seem to affect the overall pace of development of the cells.

**Figure S3.**
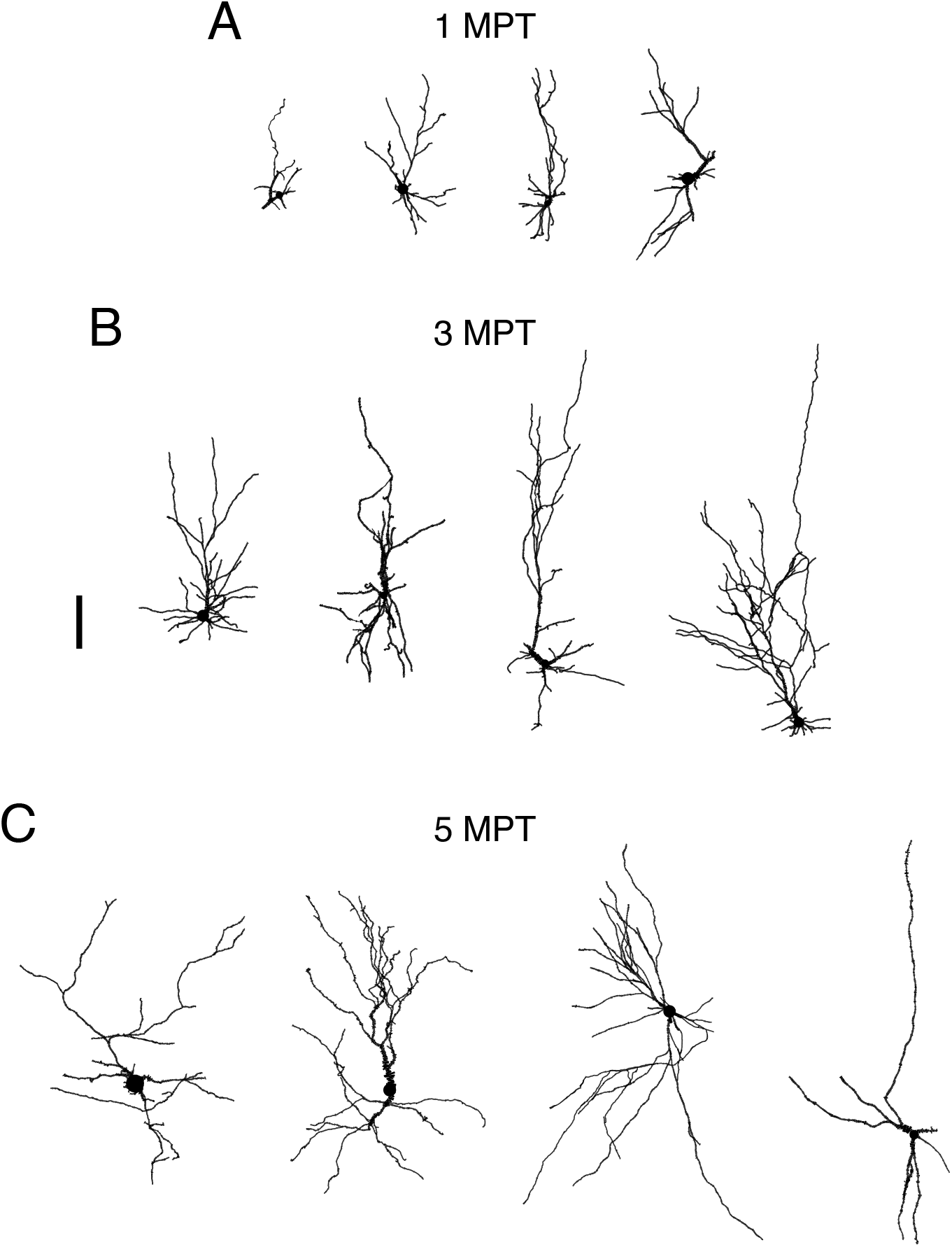
Representative human cell morphologies at various developmental stages. (A-C) Four digitally reconstructed cell morphologies at 1, 3 and 5 MPT. Notice the increase in dendritic complexity and overall length. Scale bar, 100 *µ*m.

**Figure S4.**
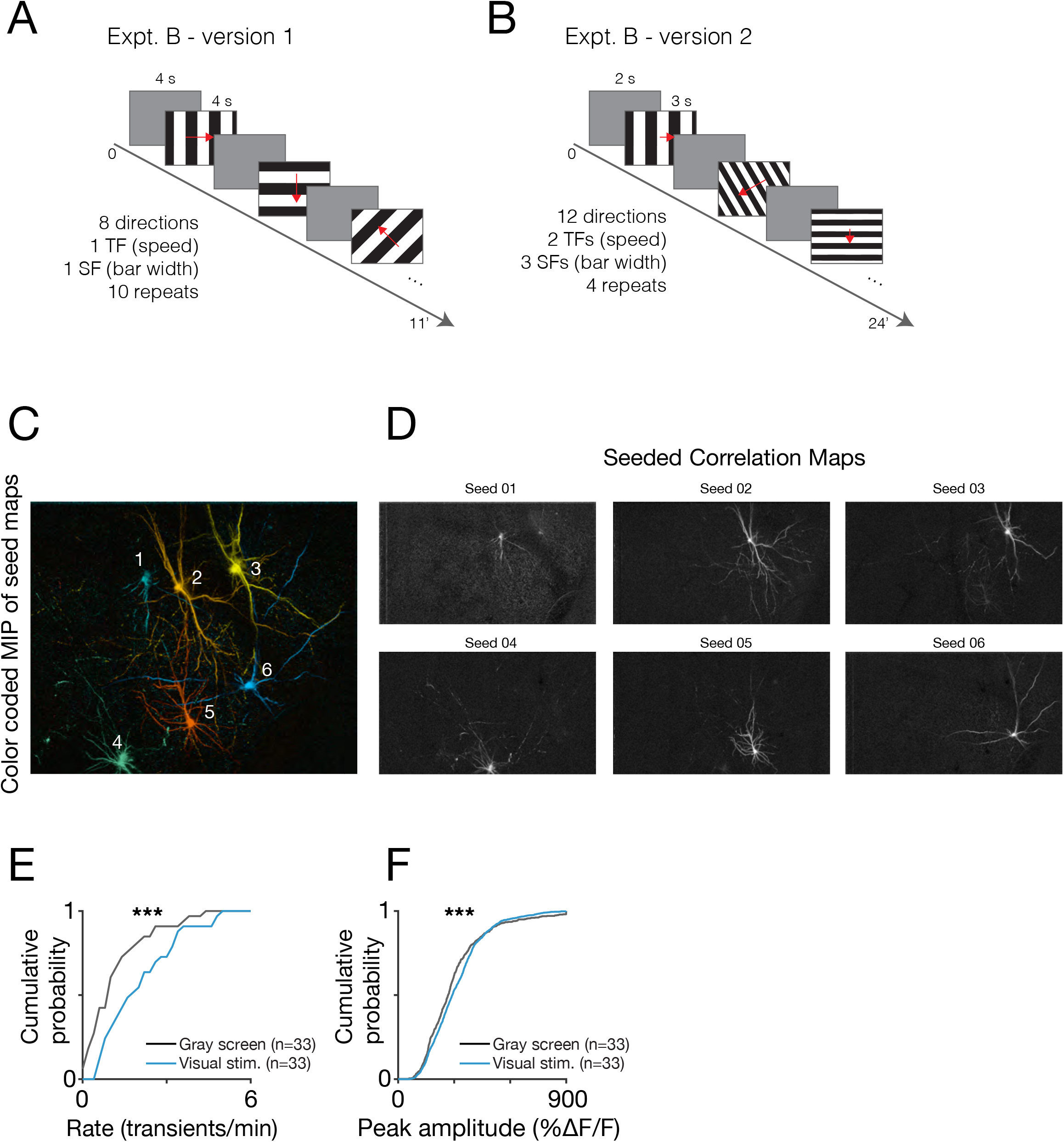
In vivo calcium imaging experimental paradigm. (A,B) Description of the visual paradigms used in this study. (A) In version 1, gratings (SF=0.05 cpd, TF=4 Hz) drifted in 8 directions. (B) In version 2, we used gratings drifting in 12 direction in combinations of two TFs (1 and 4 Hz) and three SFs (0.04, 0.08 and 0.16). (C) Maximum intensity projection of seeded correlation maps for 6 neurons. (D) Individual seeded correlation maps. To generate these maps, we correlated the average fluorescence trace of each “seed” neuron with the time course of each pixel in the movie. These correlation values are then collapsed into a single 2D image that provides a detailed image of the morphology of the seed neuron. (E,F) Cumulative distributions for transient rate and amplitude for matched cells recorded during grey screen and visual stimulation.

**Figure S5.**
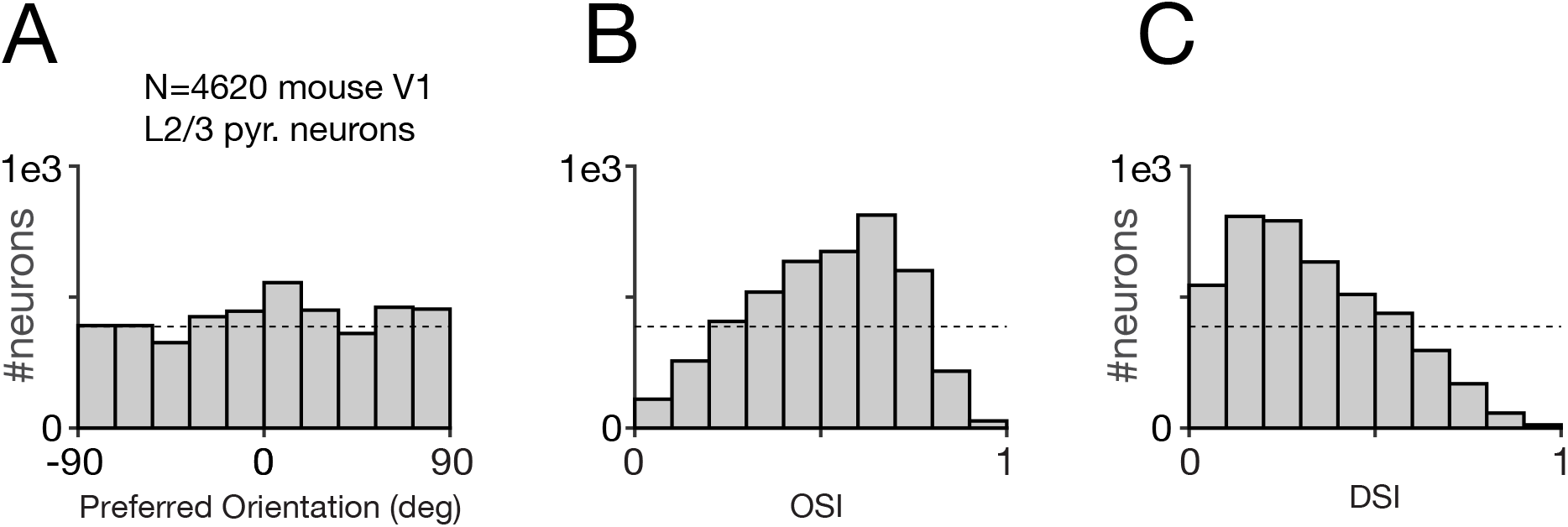
Features of mouse neurons in response to visual stimulation in vivo. (A-C) Probability distributions of 4620 mouse neurons for (A) preferred orientation, (B) orientation selectivity index and (C) direction selectivity index. These same data are presented as reference in Figure 7B-D.

**Supplementary Table S1. Neuron properties.**

See attached Excel file.

